# Full-length genome sequence of segmented RNA virus from ticks was obtained using small RNA sequencing data

**DOI:** 10.1101/2020.03.24.004796

**Authors:** Xiaofeng Xu, Jinlong Bei, Yibo Xuan, Jianyuan Chen, Defu Chen, Stephen C. Barker, Samuel Kelava, Bingjun He, Shan Gao, Ze Chen

**Affiliations:** State Key Laboratory of Veterinary Etiological Biology and Key Laboratory of Veterinary Parasitology of Gansu Province, Lanzhou Veterinary Research Institute, Chinese Academy of Agricultural Science, Lanzhou, Gansu 730046, P.R.China; Agro-Biological Gene Research Center, Guangdong Academy of Agricultural Sciences, Guangzhou, Guangdong 510640, P. R. China; Guangdong Laboratory for Lingnan Modern Agriculture, Guangzhou, Guangdong 510642, P. R. China; Hebei Key Laboratory of Animal Physiology, Biochemistry and Molecular Biology, College of Life Sciences, Hebei Normal University, Shijiazhuang, Hebei 050024, P.R.China; College of Life Sciences, Nankai University, Tianjin, Tianjin 300071, P.R.China; School of Chemistry and Molecular Biosciences, The University of Queensland, Brisbane, QLD 4072, Australia

**Keywords:** virus detection, mogiana, full-length genome, 5’ sRNA, 3’ sRNA

## Abstract

In 2014, A novel tick-borne virus of the genus *Flavivirus* was first reported from the Mogiana region in Brazil. This virus was named the Mogiana tick virus (MGTV). Later, MGTV was also named as Jingmen tick virus (JMTV), Kindia tick virus (KDTV), Guangxi tick virus (GXTV) etc. In the present study, we used small RNA sequencing (sRNA-seq) to detect viruses in ticks and detected MGTV in *Amblyomma testudinarium* ticks, which had been captured in Yunnan province of China in the year of 2016. The full-length genome sequence of a new MGTV strain Yunnan2016 (GenBank: MT080097, MT080098, MT080099 and MT080100) was obtained and recommended to be included into the NCBI RefSeq database for the future studies of MGTV. Our phylogenetic analyses showed that viruses named MGTV, JMTV, KDTV and GXTV are monophyletic: the MGTV group (lineage) of viruses. We show, for the first time, that 5′ and 3′ sRNAs can be used to obtain full-length sequences of the 5’ and 3’ ends of, but not limited to genome sequences of RNA viruses. And we proved the feasibility of using the sRNA-seq based method for the detection of viruses in a sample containing miniscule RNA.

## Introduction

Next generation sequencing (NGS) has been widely applied to virus and viroid discovery in plants and animals. Compared to other NGS based methods that require viral enrichment and concentration, the small RNA sequencing (sRNA-seq) methods simplify virus detection and indeed have several other advantages ^[1]^. The sRNA-seq based method was originally used for viral detection and identification in plants ^[2]^ and in invertebrates ^[3]^. Although the sRNA-seq based method does not perform as well in the detection of mammalian viruses as in the detection of plant or invertebrate viruses, we still detected eight mammalian viruses: HPV-18 (Human Papillomavirus type 18) ^[4]^, HBV (Hepatitis B Virus) ^[4]^, HCV (Hepatitis C Virus) ^[4]^, HIV-1 (Human Immunodeficiency Virus type 1) ^[4]^, SMRV (Squirrel Monkey Retrovirus) ^[4]^, EBV (Epstein-Barr Virus) ^[4]^, SARS-CoV (Severe Acute Respiratory Syndrome Coronavirus) ^[5]^ and a DNA segment of ASFV (African Swine Fever Virus) ^[6]^. The discovery of featured RNA fragments, including siRNA duplexes ^[7]^, 5′ and 3′ end small RNAs (5′ and 3′ sRNAs) ^[8] [9]^, palindromic small RNAs (psRNAs) and complemented palindromic small RNAs (cpsRNAs) ^[5]^, has increased our capacity to detect viruses in mammals. Moreover, we found that 5′ and 3′ sRNAs can be used to annotate nuclear non-coding and mitochondrial genes at 1 bp resolution ^[10] [11]^. Then, we found that 5′ and 3′ sRNAs can be used to obtain the full-length 5’ and 3’ ends of, but not limited to genome sequences of RNA viruses at 1-bp resolution.

Ticks transmit a great variety of infectious agents to humans and other animal species, including viruses of the Flaviviridae family among the most common tick-borne viruses ^[12]^. With the development of NGS, a number of studies have used metagenomics to search for tick-associated pathogens ^[13]^. However, the metagenomics methods using DNA can not be applied to detect RNA viruses without DNA stages. So the transcriptomics approaches using total RNA were also applied to detect tick viruses ^[14]^. The sRNA-seq based method has been successfully used in the detection of Rickettsia in ticks ^[15]^. As far as we know, there had not been previous reports of virus detection in ticks using the sRNA-seq based method until the detection of the DNA segment of ASFV ^[6]^. Although very high-depth sRNA-seq data was used to detect a DNA segment of ASFV, the coverage of the ASFV reference genome was still very low. This suggested that the sRNA-seq based method does not perform as well in the detection of DNA viruses as in the detection of RNA viruses.

In our previous study ^[6]^, we used sRNA-seq to detect viruses in ticks. Subsequent analyses of the detected viruses led to the discovery of a new strain of RNA virus. In 2014, this virus was first reported as a novel tick-borne virus of the genus *Flavivirus* from the Mogiana Region in Brazil ^[12]^. Later, this virus was also named as Jingmen tick virus (JMTV), Kindia tick virus (KDTV) or Guangxi tick virus (GXTV) in some data from the NCBI GenBank database (**Results and Discussion**). In humans, viruses closely related to MGTV were found to be primarily associated with patients in Kosovo ^[16]^. In the present study, we reported a new MGTV strain Yunnan2016 detected in *Amblyomma testudinarium* ticks ^[17]^ and also aimed to achieve the following research goals: (1) prove the feasibility of using the sRNA-seq based method in the detection of viruses in a sample (two ticks) containing miniscule RNA; (2) show how to obtain the full-length genome sequence of a RNA virus using sRNA-seq data; and (3) to provide a high-quality and well curated reference genome for the future studies of MGTV.

## Results and Discussion

### Detection of viruses in ticks using sRNA-seq data

Ticks of *Amblyomma testudinarium, D. nuttalli, D. niveus* and *D. silvarum* were captured in our previous studies. Then, the sRNA-seq data from these ticks was produced and deposited in the NCBI SRA (accession numbers SRP084097 and SRP178347; **Table 1**). Using VirusDetect (**Methods**), MGTV was detected in *A. testudinarium* ticks (SRA: SRP084097), but not in *D.nuttalli, D. niveus* nor *D. silvarum* (SRA: SRP178347) that were used as negative controls. Since the *A. testudinarium* ticks were captured in Yunnan province of China in the year of 2016, the new MGTV strain was named the Yunnan2016 strain of MGTV. As a segmented RNA virus, MGTV comprises four RNAs in its genome, two of which (RNA1 and RNA3) are related to the nonstructural protein genes of the genus Flavivirus (family Flaviviridae), whereas the other two segments (RNA2 and RNA4) are unique to MGTV. VirusDetect (**Methods**) uses the closest reference sequence to report the detected virus. The reference genome of the MGTV strain Xinjiang2016 (GenBank: MK174251, MK174244, MK174230 and MK174237) was sequenced from wild rodents captured in Xinjiang province of China. The sRNA-seq data of *A. testudinarium* (SRA: SRR4116826) covered 86.71% of the Xinjiang2016 genome with an average depth of 46.66 (**Table 1**). The sample of *A. testudinarium* had much more sRNA-seq data aligned to the Xinjiang2016 genome (**Figure 1B**) or the Yunnan2016 genome (**Figure 1A**) than the samples of three other species (**Figure 1C**). RNA1, RNA3, RNA3 and RNA4 of the MGTV strain Yunnan2016 were assembled into contigs. PCR amplification coupled with Sanger sequencing (**Methods**) was used to fill the gaps between contigs and confirm the genome assembly. 93.7 % (2879/3073) of RNA1, 90.6 % (2528/2790) of RNA2, 88.3 % (2468/2795) of RNA3 and 95.2% (2619/2752) of RNA4 were confirmed by Sanger sequences [the polyA tails of 3’-UTRs were not part of these calculations].

**Table 1.** Genome coverage and average depth of the MGTV strain Yunnan2016 “Data” (the first column) was aligned to the reference genome (the 3rd column) to obtain the information on the fourth and fifth columns. “Data” was unique for each run of sRNA-seq data in the NCBI SRA database. The virus strain Yunnan2016 (GenBank: MT080097, MT080098, MT080099 and MT080100) was detected in the present study. The reference genome of the MGTV strain Xinjiang2016 (GenBank: MK174251, MK174244, MK174230 and MK174237) was used to report Yunnan2016. “Coverage” and “Depth” indicated the genome coverage and average depth, respectively (**Methods**).

| Data | Species | Reference | Coverage | Depth |
| --- | --- | --- | --- | --- |
| SRR4116826 | *A. testudinarium* | Xinjiang2016 | 86.47% | 46.52 |
| SRR4116826 | *A. testudinarium* | Yunnan2016 | 97.37% | 91.11 |
| SSR8439389<br>SRR8439390 | *D. nuttalli* | Yunnan2016 | 18.40% | 2.58 |
| SRR8432408 | *D. niveus* | Yunnan2016 | 18.62% | 2.91 |
| SRR8432409 |  | Yunnan2016 | 20.72% | 2.94 |
| SRR811197094 | *D. silvarum* | Yunnan2016 | 8.11% | 4.22 |
| SRR811197093 |  | Yunnan2016 | 24.38% | 10.31 |

**Figure 1.**
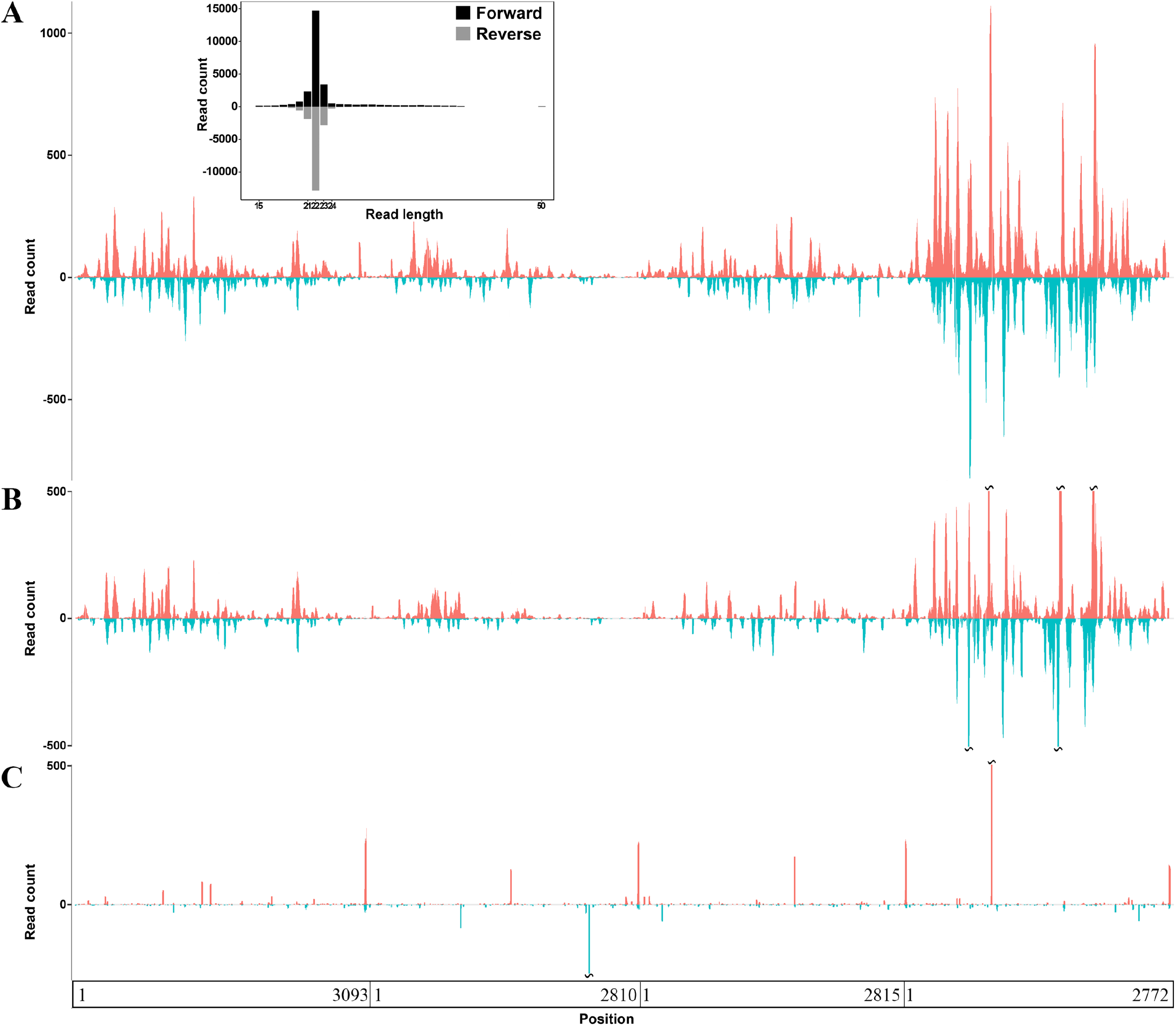
Genome coverage and average depth of the MGTV strain Yunnan2016 The y-axis represents the read-counts for each genomic position. **A.** The x-axis represents positions in the reference genome of the MGTV strain Yunnan2016 (GenBank: MT080097, MT080098, MT080099 and MT080100) and two *A. testudinarium* ticks were used to produce the sRNA-seq data **B.** The x-axis represents positions on the reference genome of the MGTV strain Xinjiang2016 (GenBank: MK174251, MK174244, MK174230 and MK174237) and two *A. testudinarium* ticks were used to produce the sRNA-seq data; **C.** The x-axis represents positions on the reference genome of the MGTV strain Yunnan2016 and three species of ticks (*Dermacentor nuttalli*, *D. niveus* and *D. silvarum*) which were used to produce the sRNA-seq data for our negative controls.

### Full-length genome sequence of the MGTV strain Yunnan2016

In our previous study, we proposed that 5’ and 3’ sRNAs can be used to annotate nuclear non-coding and mitochondrial genes at 1-bp resolution, and to identify new steady RNAs, which are usually transcribed from functional genes ^[9]^. In the present study, we used 5’ and 3’ sRNAs (**Figure 2A**) to obtain the full-length genome sequence of the new MGTV strain Yunnan2016. The 5’ ends of all RNAs in the Yunnan2016 genome had the sequence motif AG[T]_2-3_[A]_4-6_[C/G]_n_AAGTGC (**Figure 2B**), where [C/G]_n_ represents a GC-enriched region. The 3’ ends of all RNAs in the Yunnan2016 genome have an AC-enriched region (**Figure 2B**). RNA1, RNA2, RNA3 and RNA4 of Yunnan2016 with lengths of 3,093, 2810, 2815 and 2772 bp were submitted to the NCBI GenBank database under the project accession numbers MT080097, MT080098, MT080099 and MT080100, respectively). The length of the polyA tail in each 3’-UTR of these RNAs was set as 20 bp. The sRNA-seq data of *A. testudinarium* (SRA: SRR039620) covered 97.37% of the full-length genome sequence of Yunnan2016 with an average depth of 91.11 (**Table 1**). 58.5% (26668/45563) of the virus reads were aligned to RNA4 (**Figure 1B**). Although MGTV is a positive-sense single-stranded RNA (+ssRNA) virus, the proportion of sRNA-seq reads aligned to the viral positive- and negative-strand was 1.42 (26767/18796).

**Figure 2.**
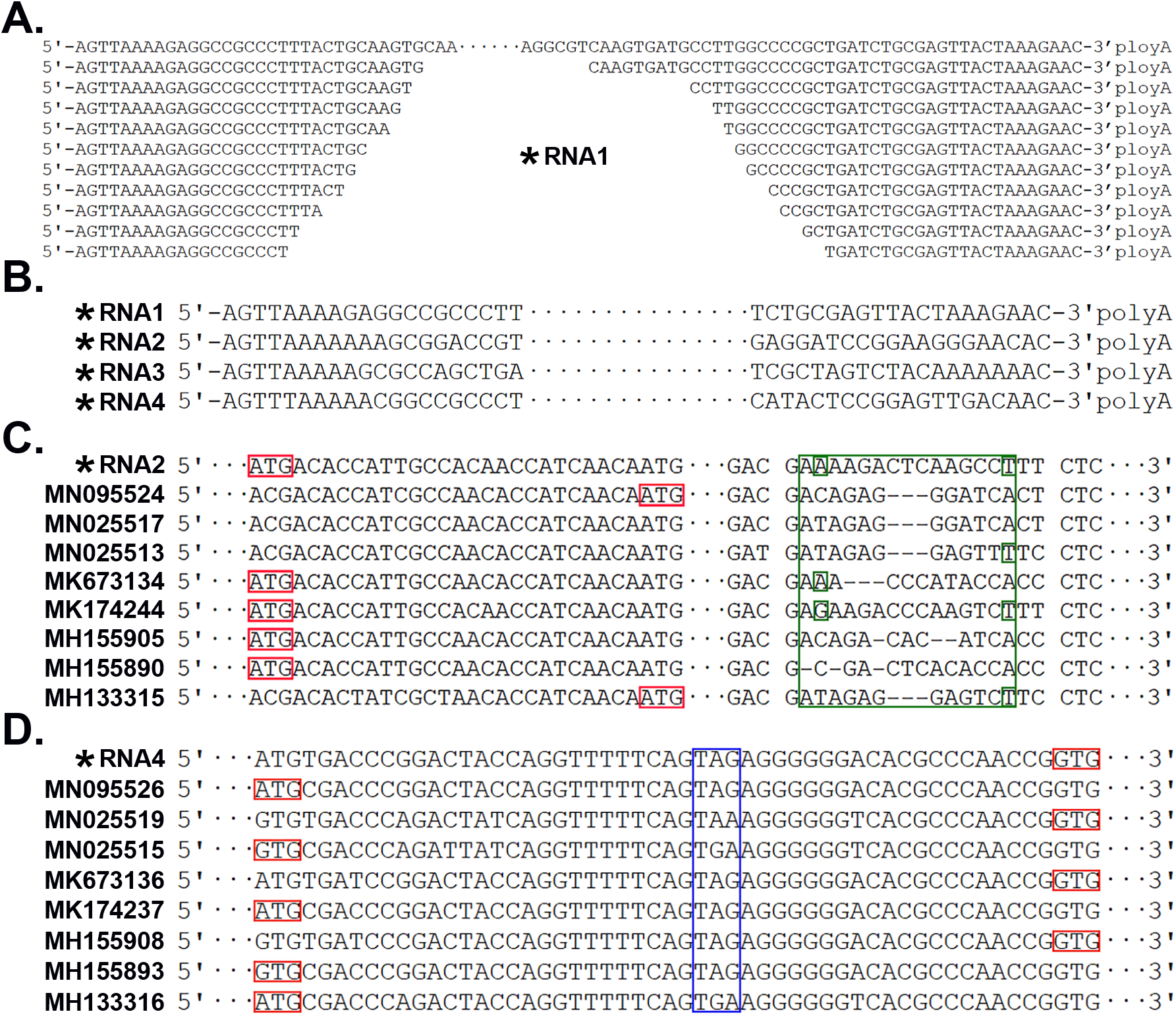
The full-length genome sequence of the MGTV strain Yunnan2016 * the MGTV strain Yunnan2016 (GenBank: MT080097, MT080098, MT080099 and MT080100). **A.** 5’ and 3’ sRNAs were used to obtain the full-length 5’ and 3’ ends of RNA1; **B.** The 5’ ends and 3’ sRNAs of all RNAs in the Yunnan2016 genome; **C.** Start codons are marked in red boxes and RNA2 has a 15- or 18-bp variable region; **C.** Start codons and stop codons are marked in red and blue boxes, respectively. For RNA4 of some viruses, “GTG” were identified as the stop codons of the second ORF and the nearby “ATG” codons are 48-bp downstream of the “GTG” codons.

Compared to the genome coverage of 86.47% (**Methods**) and average depth of 46.52 (**Methods**) when we used the Xinjiang2016 genome for reference (**Figure 1A**), genome coverage increased to 97.37% and average depth to 91.11 % when we used the Yunnan2016 genome for reference (**Figure 1B**). Comparing the full-length genome sequence of Yunnan2016 with those of 16 other MGTV complete genomes (**Methods**), we found that all of the other complete MGTV genomes did not have the correct full-length 5’ and 3’ ends. Particularly, RNA1 (GenBank: MN025516) and RNA4 (GenBank: MN025515) of the MGTV strain TT2017-2 had 56 and 48 bp additional sequences at their 5’ ends, respectively. Further analyses showed the additional sequences were identical to internal regions of the genomes. Obviously, these additional sequences had been wrongly assembled in the previous studies. Since our results showed the high quality of our full-length genome sequence of the MGTV strain Yunnan2016, we recommend that our sequencebe included in the NCBI RefSeq database for the future studies of MGTV.

### Phylogenetic analysis of MGTV complete genomes

The putative proteins encoded by RNA1 and RNA2 are the RNA dependent RNA polymerase and the glycoprotein, respectively, whereas the putative proteins encoded by RNA4 are the capsid protein and the membrane protein. The putative protein encoded by RNA3 is unknown. In total, five protein-coding genes were annotated for the genome of the MGTV strain Yunnan2016. The RNA dependent RNA polymerase from RNA1, the putative protein from RNA3 and the capsid protein from RNA4 had lengths of 915, 809 and 255 aa, respectively. The lengths of these three proteins were the same in all 17 MGTV genomes whereas those of the other two proteins (the glycoprotein from RNA2 and the membrane protein from RNA4) differed with the other MGTV genomes. The lengths of the glycoprotein from RNA2 varied due to two mutations (**Figure 2C**).: T/C mutations in the start codons shortened the Coding Sequences (CDSs) of RNA2 by 21 bp whereas a small Insertions/Deletions (InDels) shortened the CDSs by 3 bp. Theoretically, four types of glycoproteins, with lengths of 745, 746, 754 or 755 aa, would be translated from RNA2, however, a glycoprotein with 746 aa was not observed in the 17 virus genomes. Since the lengths of the membrane protein from RNA4 varied due to one mutation T/C (**Figure 2D**), two types of membrane proteins with lengths of 522 or 539 aa can be translated from RNA4.

RNA1, RNA3 and RNA4 had 2745-, 2427- and 2351-bp complete CDS 1, 3, and 4, whereas RNA2 had a 2265-bp complete CDS 2, with a 15- or 18-bp variable region removed (**Figure 2C**). Then, four CDSs could be connected into a total CDS. Using paired Pearson correlations between CDSs of 17 viruses, the degrees of evolutionary conservation are ranked as CDS 1, 2, 3 and 4 (**Figure 3A**). Five phylogenetic trees from the 2745-, 2265-, 2427-, 2351-bp and the final CDSs were built. Although these four CDSs had large differences in their degree of evolutionary conservation, the tree topologies were still congruent (**Figure 3BCDEF**). Viruses named as MGTV, JMTV, KDTV and GXTV belong to a MGTV group with two major clades. The two branches oif Clade I had the virus strains isolated from Brazil and Guinea, and the virus strains Yunnan2016 and Xinjiang2016 (**Figure 3F**). The Yunnan2016 strain was detected in *A. testudinarium* whereas the Xinjiang2016 strain was detected in wild rodents but its biological vector is still unknown. In Clade II (**Figure 3F**), the virus strain Kosovo2013 was from humans whereas the strains TT2017-1/−2 were detected in *Rhipicephalus (Boophilus) microplus*. Viruses closely related to MGTV were primarily associated with patients in Kosovo: two of the three patients had a history of tick bites ^[16]^. However, the biological vectors of the strains infecting human are still unknown. The Antilles (France) strain was detected in a pool of *R. microplus* and *mblyomma variegatum* ticks and the RT-PCR of individual ticks revealed that MGTV existed with a higher prevalence in *R. microplus* (42%) than *A. variegatum* (5%)^[18]^.

**Figure 3.**
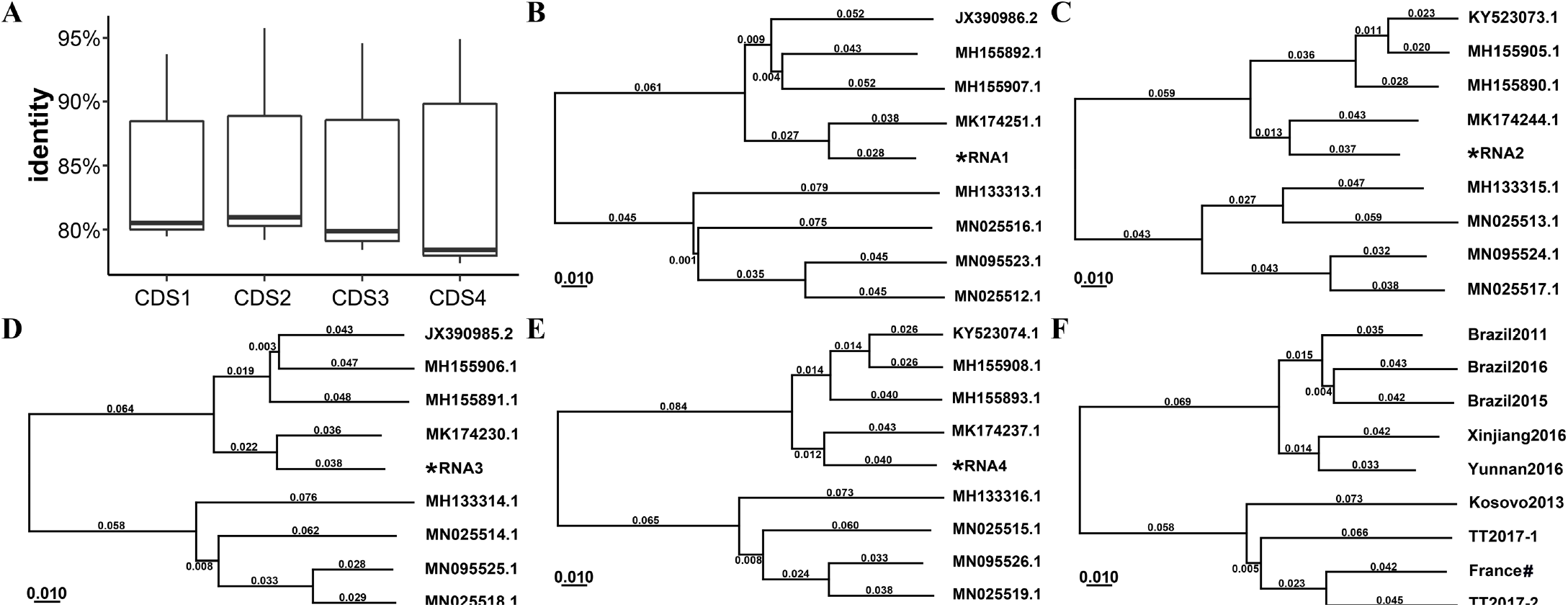
Phylogenetic analysis of Mogiana tick viruses **\*** RNA1, RNA2, RNA3 and RNA4 of the MGTV strain Yunnan2016 were submitted to the GenBank (MT080097, MT080098, MT080099 and MT080100, respectively). **A.** Paired Pearson correlations between CDSs of 17 viruses were used to account for the degrees of evolutionary conservation among four CDSs. Five phylogenetic trees were built from the 2745-bp (**B**), 2265-bp (**C**), 2427-bp (**D**), 2351-bp bp (**E**) and the final CDSs (**F**).

## Conclusions

In the present study, we reported a high-quality, fully curated and full-length genome sequence of a new strain of the Mogiana tick virus (MGTV), Yunnan2016. And we show, for the first time, that 5′ and 3′ sRNAs can be used to obtain full-length sequences of the 5’ and 3’ ends of, but not limited to genome sequences of RNA viruses. And we proved the feasibility of using the sRNA-seq based method for the detection of viruses in a sample containing miniscule RNA.

In total, 17 MGTV strains have been found in five species of ticks: *R. microplus* (9 virus strains), *R. geigyi* (1 virus strain), *A. testudinarium* (2 virus strains), *A. variegatum* (1 strain), and *A. javanense* (1 strain). MGTV strains have also been isolated from humans (3 virus strains) and rodents (1 virus strain). Among these strains, the Antilles (France) strain was detected in a pool of *R. microplus* and *A. variegatum* ticks. *R. microplus* is one of the most widely distributed tick species in the world ^[19]^. *R. microplus* primarily parasitizes livestock, but it may also feed on people in China. The detection of MGTV in *A. testudinarium* was reported for the first time in Laos. The nucleotide identity between the genome of the MGTV strain from Laos and the Yunnan2016 genome was 94.73%. Compared to other ticks, the host-range of *A. testudinarium* is very large, including mammals, amphibians, reptiles and birds, with widely distributed in tropical and subtropical areas of Asia ^[20]^. The detection of MGTV in wild rodents and humans indicates that the ticks (e.g. *R. microplus* and *A. testudinarium*) host MGTV: this hypothesis should be well investigated in future studies.

## Methods

The full-length genome sequence of the MGTV strain Yunnan2016 has been deposited into NCBI GenBank database under the project accession numbers MT080097, MT080098, MT080099 and MT080100. In the present study, the 17 MGTV complete genome sequences (including the Yunnan2016) were downloaded from the NCBI GenBank database and analysed together. One genome sequence was removed because it had too many ambiguous nucleotides. The online software CD-HIT [21] (Date: 20191212) was then used to remove redundant sequences, resulting in 9 complete genome sequences for the phylogenetic analysis. The sRNA-seq data used in the present study was also deposited in the NCBI SRA database under the project accession number SRP084097 and SRP178347 (**Table 1**). Using specific primers (**Table 2**), PCR amplification coupled with Sanger sequencing was used to fill the gaps between contigs and to confirm the assembly results.

**Table 2.**
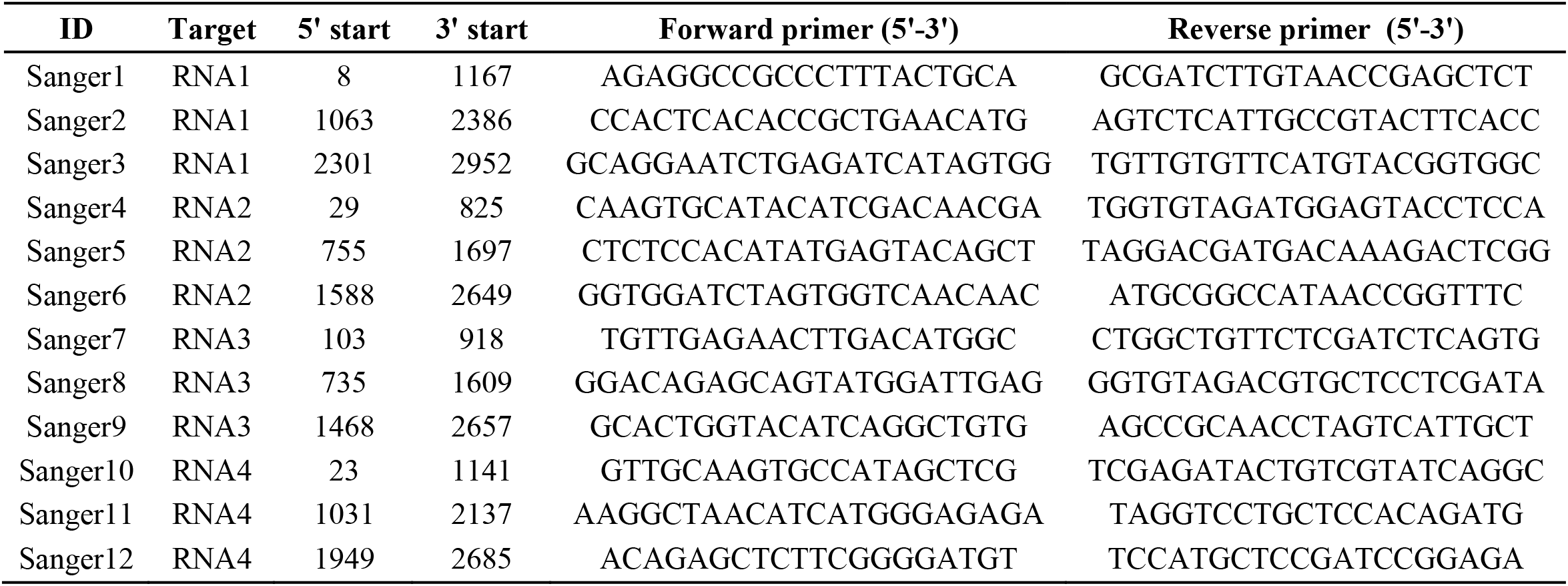
Primers for PCR amplification coupled with Sanger sequencing “ID” is unique for each Sanger sequence in the present study. RNA1, RNA2, RNA3 and RNA4 of the MGTV strain Yunnan2016 have been submitted to the NCBI GenBank database under the project accession numbers MT080097, MT080098, MT080099 and MT080100, respectively.

The software Fastq_clean ^[22]^ was used for sRNA data cleaning and quality control. The virus detection was performed using the pipeline VirusDetect [23]. For each detected virus, VirusDetect assigned a closest reference genome from the NCBI Genbank database to help characterize that virus. VirusDetect used reference genome coverage and average depth to quantify the detected viruses for verification. Genome coverage was defined as the proportion of read-covered positions divided by genome length whereas average depth was the total number of base pairs of aligned reads divided by the read-covered positions of the reference genome. Statistical computation and plotting were performed using the software R v2.15.3 with the Bioconductor packages [24].

## Acknowledgments

We appreciate the help equally from the people listed below. They are Professor Guoqing Liu, Dawei Huang, Yanqiang Liu, Associate Professor Bingjun He, Qiang Zhao, the graduate student Xiufeng Jin, Haishuo Ji from College of Life Sciences, Nankai University.

## Author contributions statements

Ze Chen and Shan Gao conceived the project and supervised this study. Shan Gao wrote the first draft of the manuscript: Stephen Barker and Samuel Kelava contributed to subsequent drafts of the manuscript and to the interpretation of the data. Xiaofeng Xu executed the experiments. Shan Gao, Jinlong Bei and Jianyuan Chen downloaded, managed and processed the data. Yibo Xuan prepared the figures and tables. Defu Chen and Bingjun He provided suggestions to this project.

